# Synchronic resisto-genesis effects identify most prevalent mechanism in *Acinetobacter baumannii*

**DOI:** 10.1101/2022.03.03.482757

**Authors:** Najmeh Alaei, Abbas Bahador, Azadeh Alaei

## Abstract

*Acinetobacter baumannii* as deadliest infection agent to ICU patients challenge experimental treatment in lack of new antibiotic and non-optimal recommendations. Despite confirm importance of infectious ICU sources, IC2 strains, ISA*ba*/*bla*_OXA_ genes and colistin in adequate concentrations or combination therapy due to potential heteroresistance; this study reveals unusual hypersensitive colistin- and sensitive aminoglycosides-resistant strains using designed methods, antibio-dendrogram classification and a computational table. Accordingly, lipopolysaccharide loss-mediated penetration and upregulated AdeAB pump-mediated compounds exclusion, are found to be causes of these abnormalities. Consequently, these simultaneous resistances termed synchronic resisto-genesis effects, drive population and create abnormal multidrug-resistant strains; while, delay extensively drug-resistant phenotype. This phenomenon predicts the predominant determinant mechanisms and makes abnormal behavior comprehensible among antibiotic-phenotypic levels or in vitro and vivo observations. Therefore, this method provides preclinical information to address therapeutic resistance in future microbial or cancer cell studies, and improves drug choosing in combination therapies, like use of inhibitory drugs.

## Introduction

The multidrug-resistance progressed *Acinetobacter baumannii* (*AcB*) in competition with *Pseudomonas aeruginosa*(Karageorgopoulos & Falagas, 2008; Kim *et al*, 2009; Vincent *et al*, 2009) to cause the most threatening infections including ventilator-associated pneumonia (VAP),(Peleg & Hooper, 2010; Wenzler *et al*, 2016) bacteremia,(Munoz-Price & Weinstein, 2008; Tabah *et al*, 2012) and Gram negative-secondary meningitis(van de Beek *et al*, 2010) in the ICUs.(Munoz-Price & Weinstein, 2008; Tabah *et al*., 2012)

Along with multidrug-resistant (MDR) infections, multiple-specificity of long-lasting survival on surfaces and global expansion highlighted molecular identification techniques of *AcB* reservoirs for preventive monitoring.(Karageorgopoulos & Falagas, 2008; Munoz-Price & Weinstein, 2008) Accordingly, from late 1990s, the sophisticated amplified fragment length polymorphism (AFLP) classification as the most robust population structure mapping method depicts standardly distribution of predominant *AcB* worldwide clonal lineages to discriminate genotypes to subspecies by whole genome analysis.(Janssen *et al*, 1996)

To date, various methods identifying clonal based on PCR and sequencing have been replaced the costly AFLP fingerprinting analysis, the most abundant of which are MLST, 3LST rep-PCR, and SBT. Although of these, the golden method, MLST in Pasteur scheme is expensive, trilocus sequence type (3LST) as a powerful screening criterion for large study populations distinguishes traditional international clones (ICs 1-3) without no sequencing by two selective multiplex PCR of three genes including endogenous *bla*_OXA-51-like_ gene evolving in *AcB*.(Zarrilli *et al*, 2013)

Resistance genes may also contribute to incidence, thereby over the last decade *AcB* outbreaks surveillance has been accompanied by tracing particularly carbapenems-resistance genes, to the extent that the WHO has given priority to carbapenems-resistant *AcB* (CR-*AcB*) treatment research.(Tacconelli *et al*, 2018) To this end, in this study the dominant role of ISA*ba*/*bla*_OXA_ genes in carbapenems-resistance were determined in contrast to other determinants comprising other β-lactamase enzymes, penicillin binding protein changes, permeability reduction and efflux pumps such as AdeABC.(Evans & Amyes, 2014; Karageorgopoulos & Falagas, 2008; Peleg & Hooper, 2010; Woodford *et al*, 2006)

The most common genes extended spectrum β-lactamases, *bla*_OXA_ encoding oxacillinase (OXA), have been already identified to 12 distinct groups based on amino acid sequences similarity, five of which are the main plasmid-borne groups in *AcB* carbapenems-resistance, apart from endogenous *bla*_OXA-51-like_ genes. The smallest genetic stimulus elements, insertion sequences (ISs) upstream of *bla*_OXA_ genes upregulate oxacillinases genes. Moreover, ISs randomly transferred additional genes such as resistance genes between bacteria by constructing composite transposons.(Evans & Amyes, 2014)

Based on these understandings, recent retrospective study with intervention approach to control the critical abundant infections with risk factors (ICU, MDR and CR-*AcB*),(Karageorgopoulos & Falagas, 2008; Munoz-Price & Weinstein, 2008; Vincent *et al*., 2009) evaluate infection dispersion, resistance trends and the highly related carbapenems-resistance genes in 100 *AcB* MDR isolates. Therefore, the new study methods were conducted here with the aim of finding infection sources and drug selection for the treatment of severe *AcB* infections in ICU patients; however, abnormal data showed more valuable results from simultaneous resistance.

## Results and Discussion

### Characterization of *AcB* Isolates of Nosocomial Infections

*AcB* in the epidemiological studies and antibiotics assays has gained recent attention due to eradication failure along with prolonged hospitalization in the ICU,(Munoz-Price & Weinstein, 2008; Tabah *et al*., 2012), creating unique multidrug-resistance, especially to carbapenems and causing the most critical nosocomial infections of VAP,(Peleg & Hooper, 2010; Wenzler *et al*., 2016) bacteremia,(Munoz-Price & Weinstein, 2008; Tabah *et al*., 2012) Gram-negative meningitis.(van de Beek *et al*., 2010)

After resistant outbreaks observation in our previous findings from the ICUs at a referral hospital of Shiraz university,(Alaei *et al*, 2016) 100 non-repetitive MDR clinical *AcB* isolates were investigated that achieved from hospitalized subjects (mean age: 39 years; 7 days to 93 years; 55(%) female, 45(%) male) and both seven specimens and locations. Predominantly samples were compiled from urinary (41%) and respiratory (30%) tracts; as well as internal (32%) and surgical (27%) ICUs (Figures 2D and 2G).

### Antibio-dendrogram and phenotypic classification

First time in our previous study, the classification method termed antibio-dendrogram in this study, was created based on antimicrobial susceptibility patterns to provide inductive method for comparing resistance information with other datasets such as genetic data.(Alaei *et al*., 2016) Considering to former role of imipenem as a broad-spectrum agent for treating *AcB* infections in ICU and the resistance increment among *AcB*s to carbapenems as the first-line therapy against Gram-negative, the evaluation of antimicrobial efficacy among the CR-*AcB* strain has gained attention.(Humphreys *et al*, 1995; Kim *et al*., 2009; Munoz-Price & Weinstein, 2008; Papp-Wallace *et al*, 2011) Therefore, antibio-dendrogram by double insertion of imipenem-resistance data introduced phenotypes that could be arranged in favour of a particular antibiotic (imipenem). Additionally, due to obtain an ambiguous and less efficient antibio-dendrogram for antibiotic-resistance diversity and rare similar antibiotic profiles, intermediate and resistant interpretive results were assumed equal (results not shown).

As a result, antibiotic profiles of 100 studied *AcBs* by antibio-dendrogram were divided into 16 distinct antibiotypes (Figure 1A). All the *AcB* isolates were completely resistant to cefotaxime, cefotaxime-clavulanate, kanamycin and amoxicillin-clavulanate; and contained resistance to at least one agent of antibiotic classes of β-lactam, β- lactamase inhibitors and third-generation cephalosporins as less effective antibiotics (83 - 100%); likewise, tigecycline, colistin, and tetracyclines as the most effective agents displayed the highest antibiotic sensitivity.

**Figure 1.**
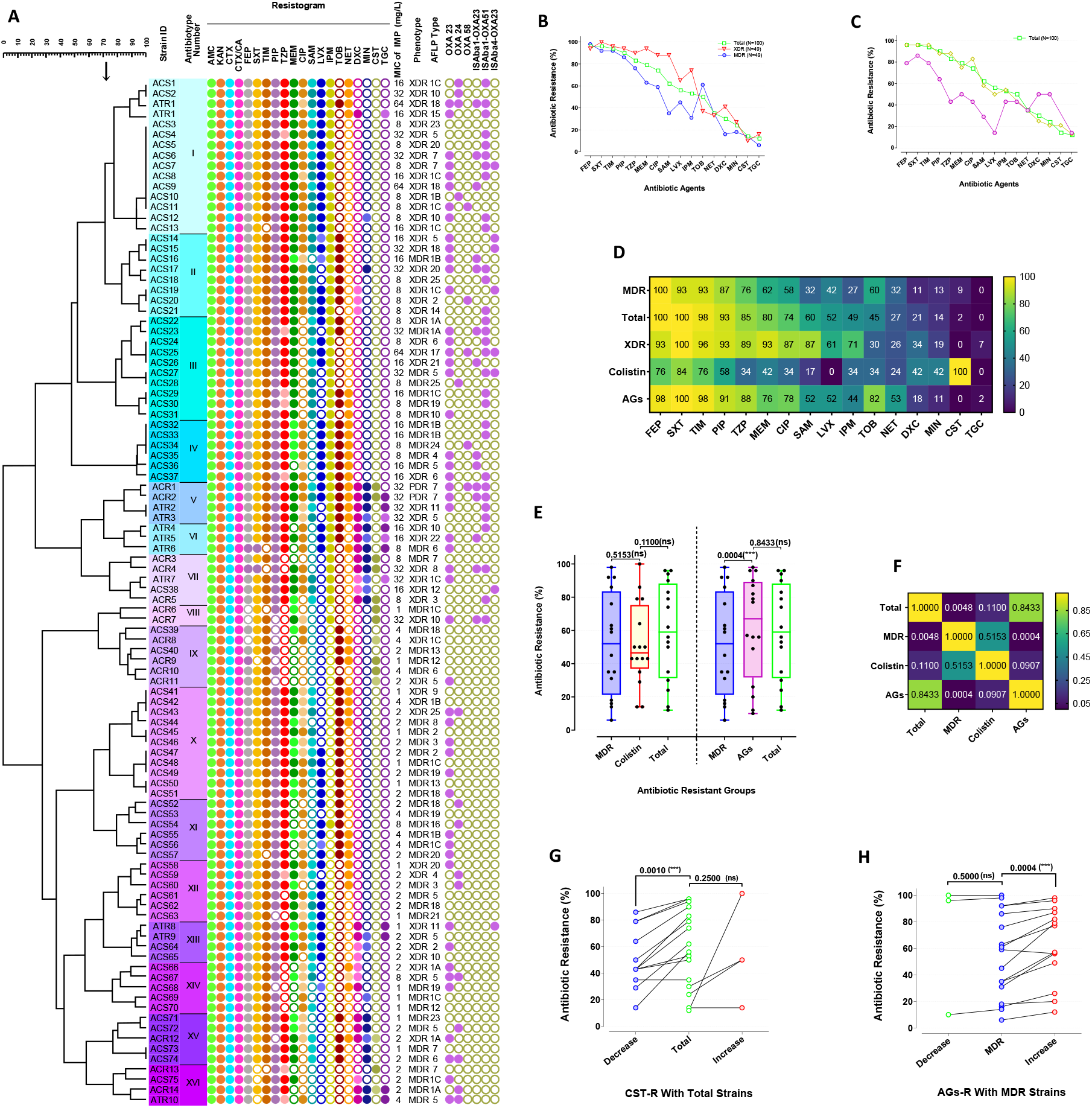
Investigation of trends in antibiotic resistance in target groups, antibiotics profiles differentiation between abnormalities. (A) Antibio-dendrogram, antimicrobial classification showing 16 antibiotypes defined with 71% similarity cut-off; antimicrobial phenotypes named based on Latin numbers; antimicrobial susceptibility patterns of 20 antibiotics; imipenem MICs (μg/mL) to compare with disk diffusion results; resistance phenotypes to determine resistance levels; AFLP genotypes to compare with antibiotic phenotypes; PCR results of genes ISA*ba*/*bla*_OXA_, respectively. Dark, light and white colours boxes denote resistance, intermediate and sensitive requests in Etest assays. Comparing the trends of antibiotic resistance percentages in total studied *AcB* strains with: (B) MDR and XDR. (C) CST-R and AGs-R. (D) Abnormalities in heatmap based on normalized values. (E) Differentiation between abnormalities distributions displayed in Box and ribbon plots and (F) Heatmap depicting statistical significance in antibiotic resistant groups. (G) Decrease and increase of antimicrobial resistances in CST-R than total strains and (H) AGs-R than MDR strains. *P values* and significance levels were determined by Wilcoxon matched-pairs signed rank test, and ns denotes not significant. Abbreviations: AGs, aminoglycosides; AMC, amoxicillin-clavulanate; CIP, ciprofloxacin; CST, colistin (polymyxin E); CTX, cefotaxime; CTX/CA, cefotaxime- clavulanate; DXC, doxycycline; FEP, cefepime; IPM, imipenem; ISs, insertion sequences; KAN, kanamycin; LVX, levofloxacin; MDR multi drug-resistant; MEM, meropenem; MIN, minocycline; NET, netilmicin; PDR, pan drug-resistance; PIP, piperacillin; SAM, ampicillin-sulbactam; SXT, trimethoprim-sulfamethoxazole; TGC, tigecycline; TIM, ticarcillin-clavulanate; TOB, tobramycin; TZP, piperacillin-tazobactam; XDR, extensively drug-resistance.

Once antibio-dendrogram is designed as the hierarchical clustering method based on frequency, represents the tree structure, such as 1, 2 and 5 antibiotypes. Nevertheless, this arrangement did not exist in some others or in several more sensitive antibiotypes colistin as the most effective drug was found unexpectedly (7-9 and 16 biotypes), as well as tobramycin and netilmicin (with the 52%, 65% effects) in 9-11 and 14 antibiotypes.

Moreover, to obtain the resistance levels three antimicrobial classifications of multi-, extensively-, and pandrug-resistant (MDR, XDR, and PDR), was defined similar to that proposed by Magiorakos *et al*.,(Magiorakos *et al*, 2012) which XDR strains were considered sensitive to be at least three classes of all nine classes, thus isolates were assigned to 49(%) MDR, 49(%) XDR and two (%) PDR strains.

### Observing Unusual Results

Thereafter, antimicrobial resistance trends were evaluated to gain access to the best treatment option at each antibiotic phenotype. Unexpectedly, computational table (Table S1) and comparing resistance trend (Figure 1B) evidenced, although XDR strains are more resistant to MDR ones, resistances to colistin and aminoglycosides (tobramycin and netilmicin) were diminished in XDR and heightened in MDR strains relative to total. Comparative analysis of resistance trends in unexpected cases illustrated that resistance percentages to all antibiotics unusually reduced except minocycline, doxycycline and tigecycline (26%, 20% and 2%, respectively) at colistin resistant (CST-R) strains as well as to most antibiotics in aminoglycoside resistant (AGs-R) strains relative to total (Figures 1C and 1D).

To determine magnitude of resistance reduction in CST-R strains, total cases without reaching statistical significance in comparison with CST-R strains by more difference than that with MDR strains, was considered the best lower criterion (Figures 1E and 1F). In this way, for abnormal CST-R (CST-R^ab^) *AcB*, the comparison with the total strains confirmed a significant decrement versus insignificant increment of resistances (Figure 1G). On the other hand, MDR strains by more difference to AGs-R strains than total was selected to explore the etiology of unexpected changes in the AGs-R strains as the lower criterion (Figures 1E and 1F), that significant increment of antimicrobial resistances was found in AGs-R strains compared to MDR strains (Figure 1H). Nevertheless, the existence of two different resistance patterns for AGs-R strains, including either sum or subscription of tobramycin and netilmicin-resistance make its interpretation confusing. For this purpose, owing to moderate aminoglycosides-resistance among studied abnormal AGs-R (AGs-R^ab^) strains, the sum of the aminoglycoside resistances indicating weak mechanisms was taken as the target antibiogram. Therefore, the antibiogram consisting of 61 tobramycin- or netilmicin-resistant cases was assumed for this phenotype that demonstrate dramatic resistance increase to all antibiotics except colistin and cephalosporins (Table S1). Crucially, abnormal resistance decrease of some agents in AGs-R strains can be due to decreased resistance of the same antibiotics or increased resistance of the opposite group. To identify abnormalities, in consistent with aminoglycosides change determined in prior computational step unexpected (Figures 1B-1D), the resistance increments were accounted abnormal.

### Explanation of Relationships in Genotypic Diversities and Carbapenems Resistance Genes

From the aspect of genetics, with prevent approach of *AcB* infections spread, two assays by different purposes were used here.

The 3LEST method to compare epidemiological genetic diversities of *AcBs* in the worldwide spatial and temporal scales, without sequencing good designed for a large scale, and cost- effectively than the PFGE and MLST identify three current main global outbreaks of sequence groups (SGs) 1-3, representing international clones (ICs) II, I and III, respectively.(Hamouda *et al*, 2010; Zarrilli *et al*., 2013) The majority of isolates (75%) were allocated to IC2/SG1 likewise 18(%) isolates to IC1/SG2, while no isolate was belonging to SG3/IC3 in our study. Others (seven isolates) without overlapping with any previous putative genotypes, were considered novel variant of SGs (SGv).

Alternatively, AFLP genotyping with high verifiability and acceptable reproducibility, closely differentiate *AcB* to intraspecific levels by the whole genome analysis relative to MLST and the amplified fragments relative to PFGE.(Hamouda *et al*., 2010; Turton *et al*, 2007; Zarrilli *et al*., 2013) Also, AFLP as a population structure study tool were modified in this study to enhance resolution. Constructed dendrogram by BioNumerics program based on 3LEST data, displayed 25 genotypes with two cut-off levels, in which eight major genotypes (1A-C, 5-7, 10 and 18) identified as the most common clinical outbreaks in this study (Figure 2A).

**Figure 2.**
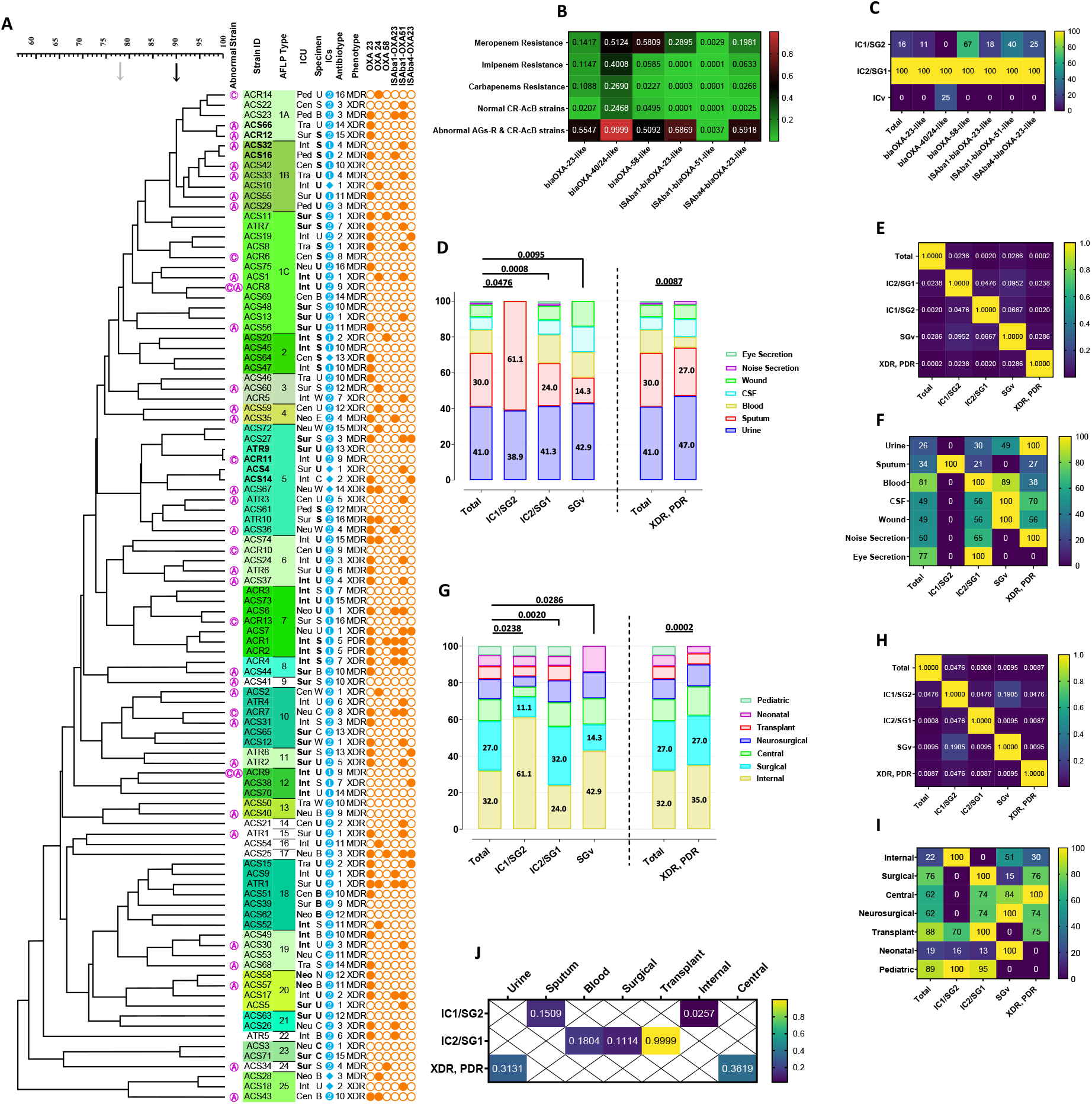
Genetic, resistance and epidemiological factors involved in *AcB* expansion. (A) Outline of abnormal AGs-R, CST-R strains; genotypic classifications of AFLP-dendrogram; clinical data to detailed transformations with respect to spatial and sample distributions; worldwide clonal lineages, another genotypes used for design AFLP-dendrogram; PCR assay of ISA*ba*/*bla*_OXA_ genes displaying role of resistance in strains distribution. Constructed AFLP-dendrogram using two cut-off levels at 78% and 90% similarity and Dice coefficient with parameter settings at one % optimization and one % tolerance limit, displayed 25 genotypes. Repeated cases are shown in boldface type and markers represent: Ⓐ, abnormal NET-R strains; Ⓒ, abnormal CST-R strains; ❷, IC2/SG1; ❶, IC1/SG2; ◆, SG variants; corresponding to the international clones (ICs) II and I. (B) Heatmap displaying significant relationships using Fisher exact tests between ISA*ba*/*bla*OXA genes and carbapenems resistances. (C) Present of ISA*ba*/*bla*_OXA_ genes in various global outbreaks, normalized to compare categories. (D-F) Comparison of genotypic and phenotypic distribution of studied *AcB* isolates in different specimens, and (G-I) ICUs. International clones of IC1 and IC2 represent SG1 and SG2, respectively, also SGv: SG variant. XDR and PDR strains indicate mot resistant samples. The Spearman’s correlation coefficients show similar distributions of infectious samples in different ICs and most resistant strains than total with respect to infectious (D and E) and spatial samples (G and H). Maximum values of worldwide clones and resistant strains were showed in F and I by input normalized data, (J) then maximum number (100) were assay to found statistical trends in this situations by Fisher exact tests. Abbreviations: B, blood; C, cerebrospinal fluid; Cen, central ICU; E, eye secretion; ICU, intensive care unit; Int, internal ICU; ISs, insertion sequences; MDR multi drug-resistant; N, nose secretion; Neo, neonatal ICU; Neu, neurosurgical ICU; PDR, pan drug-resistance; Ped, pediatric ICU; S, sputum; SG, sequence groups; Sur, surgical ICU; Tra, transplant ICU; U, urine; W, wound; XDR, extensively drug-resistance.

In terms of treatment options, resistance acquisition by either horizontal transfer or upregulation of genes through mutations or ISs induction make *AcB* challenging. Up to now, following increase usage of imipenem against *klebsiella* and environmental durability of *AcB*, *bla*_OXA_ genes as the most significant determinants involved in clinical CR-*AcB*,(Evans & Amyes, 2014; Karageorgopoulos & Falagas, 2008; Woodford *et al*., 2006) have been evolved under selective pressure into the five main oxacillinases groups *bla*_OXA-23-like_, *bla*_OXA-24/40-like_, *bla*_OXA-51-like_, *bla*_OXA-58-like_ and *bla*_OXA-143-like_, that their detection would demonstrate horizontal transfer and predominant protected contribution rather than other involved determinants. Notably, the chromosome-encoded OXA-51 weak carbapenemase and OXA-23, the most extensive acquired intermediate β-lactamases in *AcB* species, mostly upregulated with ISA*ba1* and ISA*ba4* elements and created a potentially serious problem with high resistance-mediating IS-*bla*_OXA_ structures: IS*Aba1-bla*_OXA-23-like_, IS*Aba1-bla*_OXA-51-like_, IS*Aba4-bla*_OXA-51-like_. Apart from overexpressing downstream genes, the IS elements with high prevalence could cause horizontal *bla*_OXA_s transfer between inter- or extra-species in colonized environments through genes-bracketing composite transposons.(Evans & Amyes, 2014; Higgins *et al*, 2010a)

Our observation demonstrated that among ISA*ba*/*bla*_OXA_, all studied *AcB* were positive for intrinsic *bla*_OXA-51-like_ and the *bla*_OXA-23-like_ in the 49(%) isolates routinely acquired in both carbapenems-susceptible and CR-*AcB*, similar *bla*_OXA-40/24-like_ in the 15(%) isolates. Whereas from all studied *bla*_OXA_s, *bla*_OXA-58-like_ gene in only five (%) CR-*AcB* isolates was found in unique little association with both imipenem and meropenem resistance. Moreover, The PCR data were negative for *bla*_OXA-143-like_ gene in overall. IS*Aba1* upstream of *bla*_OXA-51-like_ gene (32%) was the most widespread studied ISA*ba*/*bla*_OXA_ genes than IS*Aba1* and IS*Aba4* upstream of *bla*_OXA-23-like_ gene in 14(%) and 8(%) *AcB* isolates, respectively. Comparison of mentioned genes revealed more associations between carbapenems-resistance and all three IS genes, then imipenem-resistance was affected by ISA*ba*/*bla*_OXA_ genes. Only, the IS*Aba1-bla*_OXA-51-like_ gene with association to meropenem-resistance was the most effective determinant at carbapenems-resistance (Figure 2B). Furthermore, although relative frequency distribution of *bla*_OXA_ and IS*Aba* genes pointed roughly equal percentages in both IC1/SG2 and IC2/SG1 (Figure 2C), the *bla*_OXA-40/24-like_ gene not found significantly in any IC1/SG2 isolates (*P* = <0/0001).

### Observing Concurrency in Resistances and Etiology

Initially, colistin and aminoglycosides were thought that synergistically alter other antibiotic activities. Interestingly, published data implied that polycationic antibiotics including colistin and aminoglycosides, disrupt the outer-membrane stability through breaking down cross-bridging in competition with LPS-binding cation ions, leading to self-uptake pathway and enhanced permeability to hydrophobic-antibiotics.(Hancock, 1984; Vaara, 1992) Therefore, colistin and aminoglycosides as outer-membrane permeabilizers increased susceptibility to hydrophobic antibiotics. Nonetheless, in this study, only each Etest represents interest antibiotic effect on untreated *AcB* cells to simulate a monotherapy method.

We hypothesized that this is due to the fact that the simultaneous multi-resistance alterations conduct abnormalities in level before drug interactions, to this end similar phenotypes were followed up in the resistance mechanisms to colistin and aminoglycosides. Two main putative mechanisms modifying LPS as target site confer colistin-resistance in *AcB*: Mutation deleting LPS within the basic structural genes *lpxA,C,D*;(Moffatt *et al*, 2010) and alternatively, constitutive mutation of *pmrA, B* genes overexpressing downstream target genes adding phosphoethanolamine (Figure 3).(Adams *et al*, 2009) Notably, resistance to cationic antimicrobial peptides in some Gram-negative Enterobacteriaceae, is produced by two-component system (TCS) including *phoPQ* genes upregulated *pmrAB* genes, in which OprH protein can help with the *oprHPQ* operon cotranscription.(Macfarlane *et al*, 2000; Olaitan *et al*, 2014) Nevertheless, evolutionary elimination of PmrAB/PhoPQ cooperation is suggested in *AcB*;(Olaitan *et al*., 2014) and contrary to our findings, the PmrAB TCS induce concurrent colistin and aminoglycosides-resistance. Consequently, suggested phenotypes for AGs-R^ab^ and CST-R^ab^ strains were not in line with resultant phenotypes of simultaneous colistin- and aminoglycosides-resistance, such as synergistic effects, membrane-permeabilizer peptides, TCS activity of PmrAB or PhoPQ, and OprH protein.

**Fig. 3.**
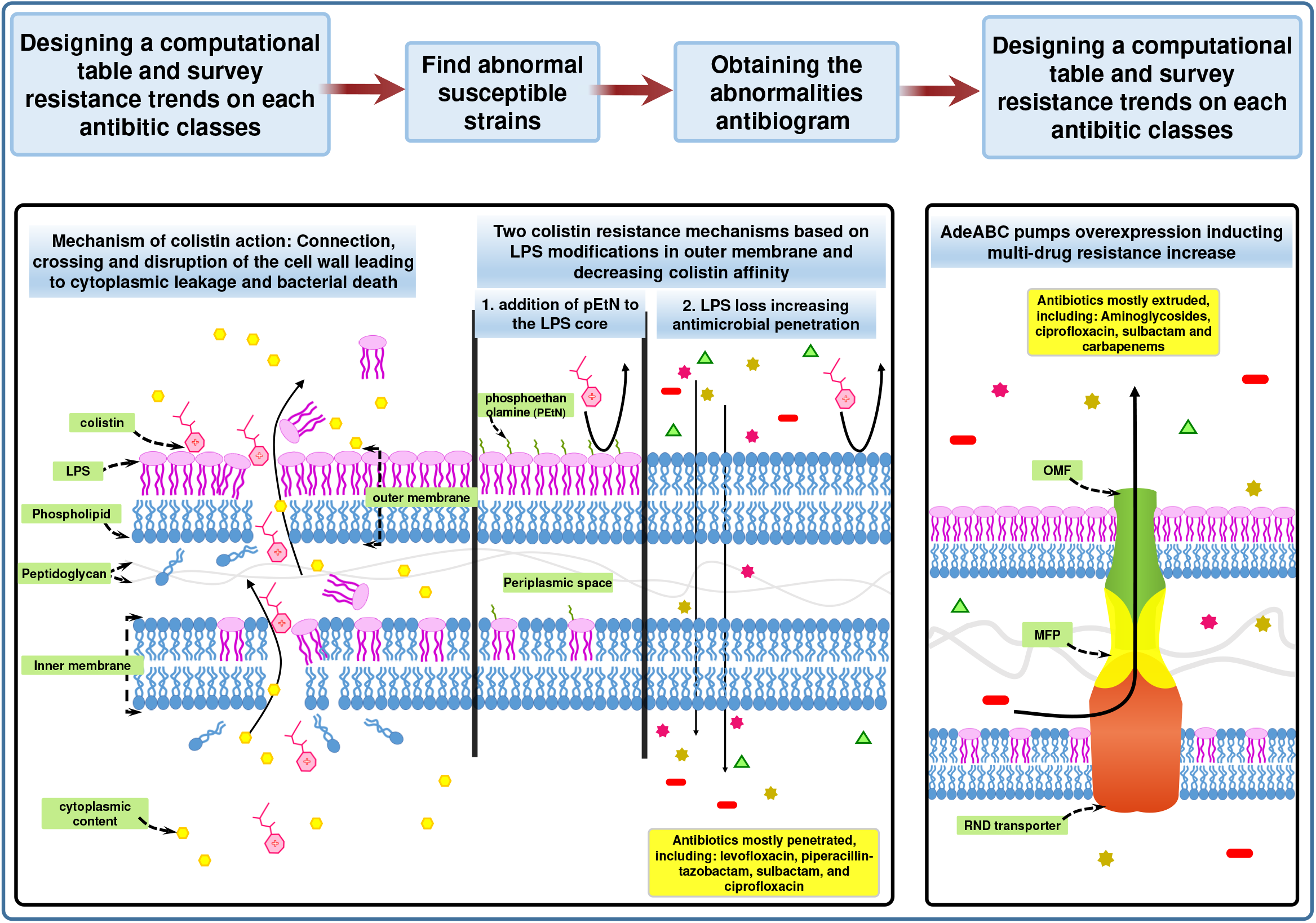
The synchronic resisto-genesis effects phenomenon identified in this study. Once detected hyper-susceptible Acinetobacter baumannii strains resistance to colistin and aminoglycosides in this study, the abnormal antibiogram results found then matched with the previously studies; consequently, the synchronic resisto-genesis effects phenomenon was introduced. In which nonspecific predominant mechanisms by moderate to mild simultaneous changes on drug-resistances (greatly here on colistin and aminoglycosides) create abnormalities, and significantly lead Acinetobacter populations as the deadliest nosocomial infections in critical ICU patients to made unpredictable experimental treatment. Two abnormalities were: Left. In colistin-resistant strains, LPS loss in outer-membrane induce colistin-resistance that increase sensitivity and penetration to all drugs, except tetracyclines and tigecycline. Right. In aminoglycoside-resistant strains, upregulated genes related to AdeABC pumps increase corresponding substrates extrude and resistance to the most antibiotics, especially aminoglycosides. Therefore, this method interestingly has beneficial effects on predicting combination therapy based on preclinical results.

The first similar observations to this sensitive CST-R^ab^ strains by Li *et al*. was reported with lowered fitness and virulence(Li *et al*, 2007) which justifies the inability to expand these strains in Figure 2A. Subsequently, Fernández-Reyes *et al*. using expression assay of similar CST-R^ab^ strains demonstrated the regulation changes of 35 proteins associated with high antibiotic susceptibility,(Fernández-Reyes *et al*, 2009) that did not consistent with the wide range of unusual changes in our study. Eventually, Moffatt *et al*. attributed this atypical permeation to *lpx* genes inactivation causing LPS loss in outer-membrane.(Moffatt *et al*., 2010) Also, numerous articles have recently confirmed substantial widely sensitivity and virulence reduction in CST-R *AcBs*(Beceiro *et al*, 2014; Durante-Mangoni *et al*, 2015; Pournaras *et al*, 2014) and more colistin efficacy in combination therapy against *AcB* compared to other Gram-negatives.(Vidaillac *et al*, 2012; Yahav *et al*, 2012)

According to this perspective, the survey of molecular parameters involved in penetration (configuration, size and distribution coefficient) showed that the complex permeability process is debatable, when the compounds have little structural difference like an antibiotic class. Thus, LPS loss-mediated colistin-resistance in *AcB* more affected on most lipophilic and hydrophilic compounds in the same antibiotic classes (Table S1). Additionally, comparing piperacillin with piperacillin-tazobactam in Table S1 referred to impressive effect of β-lactamase inhibitor when periplasmic hydrolysing enzymes are reduced upon release from defective outer-membrane.

It should be noted that despite growing evidence for CST-R^ab^ strain phenotype, the reason of tetracyclines and tigecycline-resistance increments in contrast to other antibiotics is unclear. In this respect, comparisons of estimated alterations among minocycline, doxycycline and tigecycline substantiated that hydrophobicity parameters not participate in this discrepancy because of equal log*P* values (~ −3·5), the incompatibility between molecular weights and the resistance increment magnitude as well as between differences of changes among resistances and log*D*s in three agents (Table S2). Then current study for the first time suggests that in sensitive CST-R^ab^ strains, mitigate integrity of cell-wall leads to low intracellular concentration and increased resistance to tetracyclines and tigecycline unlike others, due to high permeability of defective outer-membrane and concentration dependent activity in these three antibiotics. Therefore, in competition with amino acid carriers, tigecycline with higher affinity to ribosomes(Jenner *et al*, 2013) was less affected than other tetracyclines (Tables S1 and S2; Figures 1C, 1D and 1G).

Among the three mechanisms of aminoglycoside-resistance, the involvements of aminoglycoside-modifying enzymes with limited substrates and mutation with need to multiple binding sites of ribosome except streptomycin with a single binding site, were impossible.(Magalhães & Blanchard, 2009; Magnet & Blanchard, 2005) Eventually, developing resistance to the wide antimicrobial range specially aminoglycosides by AdeABC efflux pump with most similarity to our results was suggested as a causative factor.(Coyne *et al*, 2011; Yoon *et al*, 2015; Yoon *et al*, 2013) Efflux pumps mediating low-resistance along with severe outer-membrane barrier provide moderate-resistance at MDR *AcBs*,(Li *et al*, 2015) that with participation of other intermediate resistant mechanisms form current dissimilar AGs-R^ab^ antibiograms.(Yoon *et al*., 2015) Resistance nodulation division (RND) pump shown in Figure 3 as most extensive one of the five putative super-families, compose of three components: outer membrane factor (OMF), periplasmic membrane fusion protein (MFP), and RND transporter.(Coyne *et al*., 2011) Main intrinsic RNDs in *AcB* comprising AdeABC, AdeFGH and AdeIJK efflux pumps regulate by upstream local *adeRS* and *adeL* genes in the opposite transcriptional orientation and *adeN* repressor at 800 kbp of regulons, respectively. Overexpressing of these operons resulting from mutation in regulatory genes or the upstream ISs elements entrance, create acquired resistances by *AdeABC*, *AdeFGH* genes and intrinsic resistance by constructive expression of the *adeIJK* gene.(Li *et al*., 2015; Yoon *et al*., 2013)

In this context, literatures previously reported conflicting profiles of RND substrates. In latter study, Yoon *et al*. with superiority in using spontaneous mutants in regulatory genes or deletion of structural pump genes in non-MDR parents, precisely indicated reducing virulence and increasing resistances related to AdeABC, in which the collaboration between AdeIJK and AdeABC was in the best agreement with our findings.(Yoon *et al*., 2015) Otherwise, our observations would not be consistent without the simultaneous intrinsic resistance of AdeIJK and the acquired resistance of AdeABC to minocycline, β-lactams, and trimethoprim(Yoon *et al*., 2015) that firstly reported by Damier-Piolle *et al*. with sensitivity reduction to chloramphenicol, tetracyclines, tigecycline, macrolides, and fluoroquinolones.(Damier-Piolle *et al*, 2008) Noteworthy, except mutations or insertion of ISs in *adeRS*, both shuttle plasmid pCM88(Yoon *et al*., 2015) and BaeSR TCS(Lin *et al*, 2014) can make such situation.

In accordance with the AdeABC findings, outstanding resistance increment in AGs-R^ab^ than MDR strains in Table S1 and Figures 1C, 1D and 1H, determined cationic compounds (aminoglycosides, ciprofloxacin),(Su *et al*, 2019) sulbactam and then carbapenems(Yoon *et al*., 2015) as the preferred agents of the AdeABC pump. The RND-type multidrug transporters located at inner membrane by multiple junction sits based on molecular size and charges with regardless of PKa, captured broad-spectrum unrelated substrates from periplasm,(Colclough *et al*, 2020; Su *et al*., 2019) exception colistin, that not available in three RND due to either weight or its target in cell wall. Otherwise, like the specific channels such as TetA, they could have limited substrates.

Importantly, despite the over-resistance of abnormalities indicating AdeABC involvement, distinguish of these AGs-R^ab^ strains from normal ones was challenging. In such a case, Nemec *et al*. suggested that netilmicin as a priority for this pump and for the less likelihood of involving another mechanism, can represent overexpressing *adeABC* gene at hyper-resistant AGs-R^ab^ strains,(Nemec *et al*, 2007) denoted in Figure 2A. Furthermore, in prior study of CR-*AcB* strains upregulating AdeABC, predominant contribution of pump than enzymatic mechanisms were inferred,(Yoon *et al*., 2015) that in comparison of 18 abnormal resistant *AcBs* resistant to netilmicin and carbapenems with other CR-*AcB* strains (Figure 2B), the significant relationship reduction in between ISA*ba*/*bla*_OXA_ genes and carbapenems-resistance was in line with this results.

Consequently, this study demonstrates that LPS-loss converts XDR to MDR strains, when causes hypersensitive CST-R *AcB* or unusually MDR CST-R phenotype. While upregulated AdeABC delays promoting to the XDR phenotype in the lack of other mechanisms by reducing horizontal gene transfer and defecting biofilm formation, despite appearing MDR AGs-R strains.(Colclough *et al*., 2020) Ultimately, observed concurrent multi-resistance changes, termed the synchronic resisto-genesis effects phenomenon, occurs before drug interaction by nonspecific mechanism (permeability or efflux) that already confused with drug synergy. In fact, this computational table evident the differences of antibiogram and treatment strategies between different resistance levels (XDR and MDR) and in overall, or forecast shared multi-resistance mechanisms like adeABC to determine appropriate combination therapy.

### Choosing Appropriate Treatment and Using Synchronic Resisto-Genesis Effects Phenomenon in Treatment of Resistant Diseases

Reviving old drugs by improved formulation alike tigecycline and colistin, and optimizing combination therapy like synergistically sulbactam employment, have been suggested, although administration recommendations against MDR *AcB* strains (*AcBs*) are still controversial.(Karageorgopoulos & Falagas, 2008; Munoz-Price & Weinstein, 2008; Penwell *et al*, 2015)

The suitable agents here were generally colistin, tigecycline, tetracyclines and aminoglycosides, while high sensitivity to imipenem, levofloxacin and sulbactam in MDR, CST-R and AGs-R *AcBs* were remarkable (Figure 1D). These relationships are also inversely found in reverse as shown in Table S1, thus colistin is the best treatment in all resistant groups except MDR strains, while synchronicity with aminoglycosides resistance is negligible. It should be noted that for local experimental therapy based on treatment strategies, in referral hospital the XDR strains is more expected; contrarily, in small hospitals by less-resistance the MDR strategy is suggested. Similarly, Table S1 attested that resistant strains to both carbapenems were highly resistant to sulbactam, also all imipenem-resistant *AcBs* were sensitive to meropenem and sulbactam, and tigecycline co-occurred by doxycycline-resistance. Therefore, this phenomenon without any complex molecular technique disclosed the resistances concurrent and shared mechanism between carbapenems and sulbactam, or between doxycycline and tigecycline as well as unique mechanism of imipenem, which had been unknown in the past reports. In this way, in addition to discuss in stable resistance detectable at Etests,(Adams *et al*., 2009; Olaitan *et al*., 2014) the new method could recommend appropriate drug in confronting with specific resistance.

Crucially, apart from *lpx* mutants found in this study, PmrAB in 90% *AcB* strains exposed to colistin reversibly confers heteroresistance.(Lim *et al*, 2010) Nonetheless, this study recommends colistin as best choice for *AcBs* treatment in adequate doses or combination therapy such as co-administering with drug inhibitors of efflux pump or TCS PmrAB to prevent risk of resistance evolution under antibiotic selective pressure.(De Silva & Kumar, 2019; Li *et al*., 2015) Notably, TCSs adapt Gram-negatives in response to environmental stimuli and broadly contribute to survival and virulence as shown in here for *AcB*. In this respect, dominant TCSs in the current study vary organism behaviour in vitro and vivo, since either AdeRS regulating adeABC or PmrAB in divalent cations, acidic pH and Fe3+ concentrations, by LPS modifications induce resistance to aminoglycosides and colistin (cationic peptides), respectively.(Mitrophanov *et al*, 2008; Winfield & Groisman, 2004)

### Resistance factors and cross transmission implicated in distribution

Epidemiological AFLP investigation detailed retrospectively clonal evolution and spatial or infectious distribution to implement appropriate interventional controls. Tracking 100 MDR outbreaks *AcB*s from university hospital revealed main outbreak sources of respiratory and urinary tracts in accordance with prior publications(Fournier *et al*, 2006; Peleg & Hooper, 2010) and internal and surgery wards that in Figure 2A, effective control of critical *AcB* infections, especially PDR strains is good visible. Furthermore, the adaptability attributed to the most prevalent IC2 global clone(Zarrilli *et al*., 2013) with superiority in acquired resistance genes containing all ISA*ba*/*bla*_OXA_ genes was referable here in its prominent presence among AGs-R^ab^ strains (80%), which facilitated transfer in the main outbreak sources. Comparison of genotypic and phenotypic diversions in studied *AcB* showed similar percentage distribution of different specimens and locations (Figures 2D, 2E, 2G and 2H), so that dispersion of dominant clone IC1/SG2 was in good agreement with total. Only presence of IC1 in urine and sputum samples and in parallel at the internal ICU without resistance superiority in IC genotypes highlight cross transmission in referral hospital of Shiraz with environmental pollution in IC1 (Figures 2E and 2H). Nevertheless, these prominent presences of different clonal lineages (Figures 2D, 2F, 2G and 2I) were not significant, except IC1/SG2 that tended to dispense in internal ICU (Figure 2J).

As mentioned above, the expansion of AFLP clusters 1C, 10 and 7 indicated implications of XDR/PDR phenotypes and ISA*ba*/*bla*_OXA_ genes-containing strains, respectively; and pointed to resistance effect on cause of dangerous outbreaks (Figures 2A). However, the predominant existence of XDR and PDR strains representing highest resistance levels in urinary infections and central ICU in Namazi hospital, no significant difference in the presence or absence of XDR and PDR *AcB* was detected (Figure 2J). In addition, the presence of acquired ISA*ba*/*bla*_OXA_ genes in all AFLP clusters emphasised on horizontal transfer and evolution of these genes in selective pressure likewise, various clusters and seven singletons belonging most prevalent global clone of IC2/SG1 on ongoing expansion and genetic diversity.(Zarrilli *et al*., 2013).

In contrast, no prevalence of resistant cases (urine samples and central ICU as shown in Figures 2F and 2I), phenotypic dissimilarity in AFLP genotypes even among 98% similar strains and existing repeated cases among ICUs and specimens (bolded in Figure 2A), evidenced role of environmental then staff pollution before acquired resistance like gene transfer in prevalence.

Moreover, in current large-scale AFLP-dendrogram, denoted abnormalities indicate CST-R^ab^ strains presence in upper AFLP clusters in contrast to the throughout distribution of AGs-R^ab^ strains that represent role of mutations depending proliferation in CST-R^ab^ strains and of random ISs insertion more than mutations in AGs-R^ab^ strains.

## Conclusion

In conclusion, this investigation noted treatment need for the threatening MDR outbreaks by the factors of *AcB* distribution encompassing compatibility under selective pressure, environmental contamination and antimicrobial resistance, respectively. This report for the first time by terming the synchronic resisto-genesis effects phenomenon, revealed population drivers in the new mathematical method displaying unusual changes and could predict concurrent antibiotic-resistance and same unknown resistance mechanisms to apply in combination therapy also at other organisms or cancer cells for future. Therefore, this phenomenon by investigation in preclinical data of microorganisms expands the informative pharmacokinetic sources and approves use of inhibitory drugs to counteract the predominant determining mechanisms in the absence of new drugs.

## Materials and Methods

### Isolates

This retrospective study was commenced following the significant resistance observations at our previous study in 2017.(Alaei *et al*., 2016) Only compiled MDR *AcB* from different ICU wards of Namazi Hospital Shiraz Iran (Figure 1) were identified to species level using API-20NE system (bioMérieux, Marcy-l’Etoile, France), later confirmed by *gyr*B-multiplex PCR in genomics species with interpreting of Higgins *et al*.(Higgins *et al*, 2010b)

### Antimicrobial assays

In accordance with CLSI guideline,(Patel, 2017) disk diffusion method to assay all twenty antibiotic susceptibilities listed in Figure 2 (from Mast Diagnostics Ltd, Bootle, UK) and broth microdilution MIC of colistin and imipenem (assessing role of oxacillinase in carbapenems-resistance) were performed, except for tigecycline that EUCAST proposed breakpoints for *Enterobacteriaceae* were scored.(Testing & Testing, 2019) Ensuring accuracy of each test was assayed by *Escherichia coli* ATCC 25922 and *Pseudomonas. aeruginosa* ATCC 27853 as quality control organisms.

Clustering antibio-dendrogram by BioNumerics software ver.7.1 (Applied Maths, St-Martens-Latem, Belgium) was designed according to ward clustering method. For the slight resemblance between susceptibility profiles of isolates in this study, initially an ambiguous and less efficient antibio-dendrogram was obtained (results not shown). Therefore, to obtain the appropriate sub-clusters with analysable and applicable information, intermediate and resistant interpretive results were assumed equal. Moreover, in order to investigate imipenem-resistant strains, results were adjusted based on resistance to imipenem by double insertion of related data.

### Genotyping assays

Primary genetic classification by two multiplex PCR of *ompA, csuE* and *bla_OXA-51-like_* genes without sequencing as latter described in 3LST method(Turton *et al*., 2007) identified the global most common clonal lineages by three SGs representing ICs.(Zarrilli *et al*., 2013)

Developed AFLP genomic fingerprinting analysis has previously been described to indicate incidences in incidence infection sources.(Bahador *et al*, 2013) In which, once double digestion with *Mbo*I and *Mse*I enzymes (Fermentas, Lithuania) and ligation by T4 DNA ligase (Takara Bio), PCRs by limited adaptors and primers were conducted (all used adaptors and PCR primers listed in Table S4).

The final data were interpreted using BioNumerics ver.7.1 (Applied Maths, St-Martens-Latem, Belgium). Similarity levels of AFLP profiles were calculated by the unweighted pair-group (UPGMA) method.

### Molecular characterization of class D carbapenemases encoding genes and upstream ISs

Prevalence of ISA*ba*/*bla*_OXA_ genes was investigated in studied *AcB*s to distinguish carbapenems-resistance determinants. The Ambler Class D carbapenemase genes (*bla*_OXA-23, 24/40, 51, 58, 143-like_ genes), IS*Aba1* and IS*Aba4* elements upstream of the *bla*_OXA-23-like_ gene, and IS*Aba1* upstream of the *bla*_OXA-51-like_ gene, were detected, which gene sequences with their references were given in Table S4.(Bahador *et al*, 2015; Higgins *et al*., 2010a; Woodford *et al*., 2006)

### Statistical analysis

All statistical analyses were performed by GraphPad Prism 9 (Software, La Jolla, CA). For each of antibiotic-resistance trends, Wilcoxon signed rank test were used to assess significant differences between separate paired nonparametric groups (total, MDR, XDR, CST-R, AGs-R strains and increase/decrease magnitudes). Variable importance of genotypic and phenotypic distributions among specimens and locations were displayed by Spearman’s signed rank test to compare how each classified groups performed. Fisher exact tests calculated correlation between carbapenems-resistance with ISA*ba*/*bla*_OXA_ genes in groups that estimate their roles on the relevant resistances; likewise, the chi-square test to differentiate the distribution of ISA*ba*/*bla*_OXA_ genes in various ICs.

## Acknowledgements

We thank all colleagues at Shiraz Namazi Hospital who generously provided strains.

## SUPPORTING INFORMATION

**Table S1. Applied antibiotics trends in different antibiotics levels and classes, calculated physicochemical properties of antibiotics.**

**Table S2. Distribution of resistance percentages of *bla*_OXA_ and IS genes in different international clones.**

**Table S3. Summary of the target genes and adaptors sequences utilized in this study.**

